# δ-containing GABA_A_ receptors on parvalbumin interneurons modulate neuronal excitability and network dynamics in the mouse medial prefrontal cortex

**DOI:** 10.1101/2024.06.14.599033

**Authors:** Xinguo Lu, Hong-Jin Shu, Peter M. Lambert, Ann Benz, Charles F. Zorumski, Steven Mennerick

## Abstract

In medial prefrontal cortex (mPFC), fast-spiking parvalbumin (PV) interneurons regulate excitability and microcircuit oscillatory activity important for cognition. Although PV interneurons inhibit pyramidal neurons, they themselves express δ subunits of GABAA receptors important for slow inhibition. However, the specific contribution of δ-containing GABAA receptors to the function of PV interneurons in mPFC is unclear. We explored cellular, synaptic, and local-circuit activity in PV interneurons and pyramidal neurons in mouse mPFC after selectively deleting δ subunits in PV interneurons (cKO mice). In current-clamp recordings, cKO PV interneurons exhibited a higher frequency of action potentials and higher input resistance than wild type (WT) PV interneurons. Picrotoxin increased firing and GABA decreased firing in WT PV interneurons but not in cKO PV interneurons. The δ-preferring agonist THIP reduced spontaneous inhibitory postsynaptic currents in WT pyramidal neurons but not in cKO pyramidal neurons. In WT slices, depolarizing the network with 400 nM kainate increased firing of pyramidal neurons but had little effect on PV interneuron firing. By contrast, in cKO slices kainate recruited PV interneurons at the expense of pyramidal neurons. At the population level, kainate induced broadband increases in local field potentials in WT but not cKO slices. These results on cells and the network can be understood through increased excitability of cKO PV interneurons. In summary, our study demonstrates that δ-containing GABAA receptors in mPFC PV interneurons play a crucial role in regulating their excitability and the phasic inhibition of pyramidal neurons, elucidating intricate mechanisms governing cortical circuitry.

**Significance statement:** By selectively deleting δ-containing GABAA receptors in PV interneurons, we demonstrate the importance of these receptors on PV interneuron excitability, synaptic inhibition of pyramidal neurons, and circuit function.

## Introduction

The medial prefrontal cortex (mPFC) participates in higher-order cognitive functions, including decision-making, working memory, attention, and behavioral flexibility (Etkin et al., 2011; Jobson et al., 2021). Dysfunction of the mPFC has been implicated in various neuropsychiatric disorders, including schizophrenia, depression, and addiction (Shin et al., 2006; Xu et al., 2019; Jacobs and Moghaddam, 2021). This underscores the importance of understanding the neural circuits and mechanisms underlying mPFC function. Among the cell types in mPFC, parvalbumin (PV) interneurons, a subclass of GABAergic neurons found throughout the cerebral cortex, are essential in regulating the activity and network dynamics of the mPFC (Kim et al., 2016; Yang et al., 2021; Binette et al., 2023; Brady et al., 2023; Ferguson et al., 2023).

PV interneurons in the mPFC provide powerful inhibitory inputs onto the soma and proximal dendrites of pyramidal neurons, the main excitatory output neurons of the cortex. This perisomatic inhibition is essential for regulating the firing rate and timing of action potentials in pyramidal neurons, thereby gating the flow of information within the mPFC and between the mPFC and other brain regions (Sohal et al., 2009; Kvitsiani et al., 2013; Courtin et al., 2014). Additionally, PV interneurons are primarily involved in the generation and synchronization of gamma-frequency brain oscillations in the mPFC (Sohal et al., 2009; Carlén et al., 2012; Lambert et al., 2024). These high-frequency oscillations are thought to play a crucial role in coordinating neuronal activity and facilitating communication between different brain regions during cognitive processes.

Interestingly, PV interneurons themselves express GABAA receptors and receive GABAergic inhibition from other interneurons (Pfeffer et al., 2013; Pelkey et al., 2017). GABAA receptors are heteropentameric ion channels, with subunit combinations that confer distinct functional and pharmacological properties (Sigel and Steinmann, 2012). Among these, the δ subunit is of interest because δ-containing GABAA receptors are selectively localized to extrasynaptic regions and mediate tonic inhibition (Brown et al., 2002; Wei et al., 2003; Glykys et al., 2008), as well as slow phasic inhibition (Herd et al., 2013; Ye et al., 2013; Sun et al., 2018; Shu et al., 2021). These receptors are expressed by only select neuronal types and are combined with specific subunits for distinct functions. In principal neurons, δ subunits preferentially pair with α4 (forebrain) or with α6 (cerebellum) subunits. However, in interneurons, δ subunits pair with α1 subunits, resulting in distinct receptor functional properties (Milenkovic et al., 2013).

Previous studies have shown that δ-containing GABAA receptors are expressed in PV interneurons and contribute to tonic inhibition as well as high frequency oscillations in hippocampus and basal amygdala (Ferando and Mody, 2013; Yu et al., 2013; Antonoudiou et al., 2022; Lu et al., 2023). However, in mPFC, the specific role of δ- containing GABAA receptors in regulating the excitability and output of PV interneurons, and consequently their influence on pyramidal neuron activity and network oscillations, remains largely unknown.

In this study, we selectively deleted δ-containing GABAA receptors in PV interneurons using a conditional knockout (cKO) mouse model. Through electrophysiological recordings, pharmacological manipulations, and gene expression analysis, we directly investigated the impact of these receptors on the intrinsic properties and synaptic output of PV interneurons. This approach allowed us to dissect the contribution of inhibition mediated by δ-containing GABAA receptors to the overall excitability and function of PV interneurons and principal neurons within the mPFC microcircuitry. We find evidence that δ-containing GABAA receptors are important for basal excitability of PV interneurons, which checks the excitability of pyramidal neurons across a broad spectrum of oscillatory activity in response to stimulation.

## Materials and Methods

### Mice

Male and female, Ai14::PVCre (Ai14, Jackson lab, #007914; PVCre, Jackson Lab, #017320), PVCre, and *Gabrd* floxed::PVCre (Lee and Maguire, 2013) mice from 4 – 8 weeks old were used. For some experiments in which only WT PV interneurons were recorded, we used Ai14::PVCre reporter mice. Alternatively, in experiments with conditional *Gabrd* deletion (cKO), pAAV-FLEX-GFP (Addgene, #28304) was injected by retro-orbital sinus injection to label PV cells in *Gabrd* floxed::PVCre mice or PVCre littermates (Lu et al., 2023).

### Slice preparation

In accordance with protocol (22-0344) approved by the Institutional Animal Care and Use Committee at Washington University, mice were subjected to isoflurane anesthesia and subsequently decapitated. A Leica VT1200 vibratome was employed to prepare 300-μm thick coronal brain slices. During the slicing procedure, the slices were kept in an ice-cold, modified NMDG-HEPES recovery artificial cerebrospinal fluid (aCSF) solution containing (in mM): 92 NMDG, 2.5 KCl, 1.25 NaH2PO4, 30 NaHCO3, 20

HEPES, 25 glucose, 2 thiourea, 5 Na-ascorbate, 3 Na-pyruvate, 0.5 CaCl2, and 10 MgSO4 (300 mOsm; pH 7.3–7.4). Following slicing, the slices recovered in the modified NMDG-HEPES recovery aCSF at 32°C. A Na^+^-rich spike-in solution (4 ml, 2 M) was added to the recovery aCSF to gradually increase Na^+^ concentration, which improved recording success rate (Ting et al., 2018). After the recovery period, the slices were maintained in a modified HEPES holding aCSF (composition in mM: 92 NaCl, 2.5 KCl,

1.25 NaH2PO4, 30 NaHCO3, 20 HEPES, 25 glucose, 2 thiourea, 5 Na-ascorbate, 3 Na- pyruvate, 2 CaCl2, and 2 MgSO4; 300 mOsm; pH 7.3–7.4) for a minimum of 1 hour at 25°C prior to conducting experimental recordings. All drugs were sourced from Sigma-Aldrich, unless otherwise specified.

### Whole-cell patch-clamp recording

For the duration of the recording, slices were placed in a recording chamber and continuously perfused (2 ml/min, 32°C) with oxygenated, regular aCSF containing (in mM): 125 NaCl, 25 glucose, 25 NaHCO3, 2.5 KCl, 1.25 NaH2PO4, 2 CaCl2, and 1 MgCl2 (310 mOsM), equilibrated with a mixture of 95% oxygen and 5% CO2. In mPFC layer 2/3, target cells for somatic, whole-cell recording were visualized and identified using IR-DIC microscopy, which was performed with a Nikon FN1 microscope equipped with a Photometrics Prime camera. Whole-cell recordings were obtained using borosilicate glass pipettes (World Precision Instruments) with an open tip resistance ranging from 3 to 6 MΩ. The recordings were conducted using a MultiClamp 700B amplifier (Molecular Devices), a Digidata 1550 16-bit A/D converter, and pClamp 10.4 software (Molecular Devices). After the initial break-in, a 5-minute stabilization period was observed before commencing the recordings.

### Measurement of action potentials

To capture action potentials, neurons were recorded in current-clamp mode using pipettes containing a K-gluconate internal solution. This solution was composed of (in mM): 140 K-gluconate, 4 MgCl2, 10 HEPES, 0.4 EGTA, 4 MgATP, 0.3 NaGTP, and 10 phosphocreatine, with osmolarity of 290 mOsm and pH adjusted to 7.25 using KOH. To assess membrane properties and evoke action potentials, step currents starting from −50 pA were injected for 500 ms, with increments of 50 pA. To confirm the involvement of GABAARs, the non-competitive antagonist picrotoxin (PTX, 100 µM) or agonist GABA (5 µM) was bath applied for 5 min. All current-clamp recordings were done in episodic mode at 50 kHz sampling rate and 10 kHz filter cutoff frequency using an eight-pole Bessel filter.

### Measurement of spontaneous inhibitory and excitatory postsynaptic currents (sIPSCs and sEPSCs)

Phasic currents were recorded in voltage-clamp mode using pipettes filled with Cs-methanesulfonate (in mM: 130 Cs-methanesulfonate, 4 NaCl, 0.5 CaCl2, 10 HEPES, 5

EGTA, 5 QX-314, 0.5 NaGTP, and 2 MgATP; pH was adjusted to 7.3 with CsOH; 290 mOsm). To record isolated sIPSCs, neurons were held at 0 mV (see Fig. 3). To record sIPSCs and sEPSCs simultaneously, neurons were held at -40 mV (see extended Fig. 3-1).

**Figure 1.**
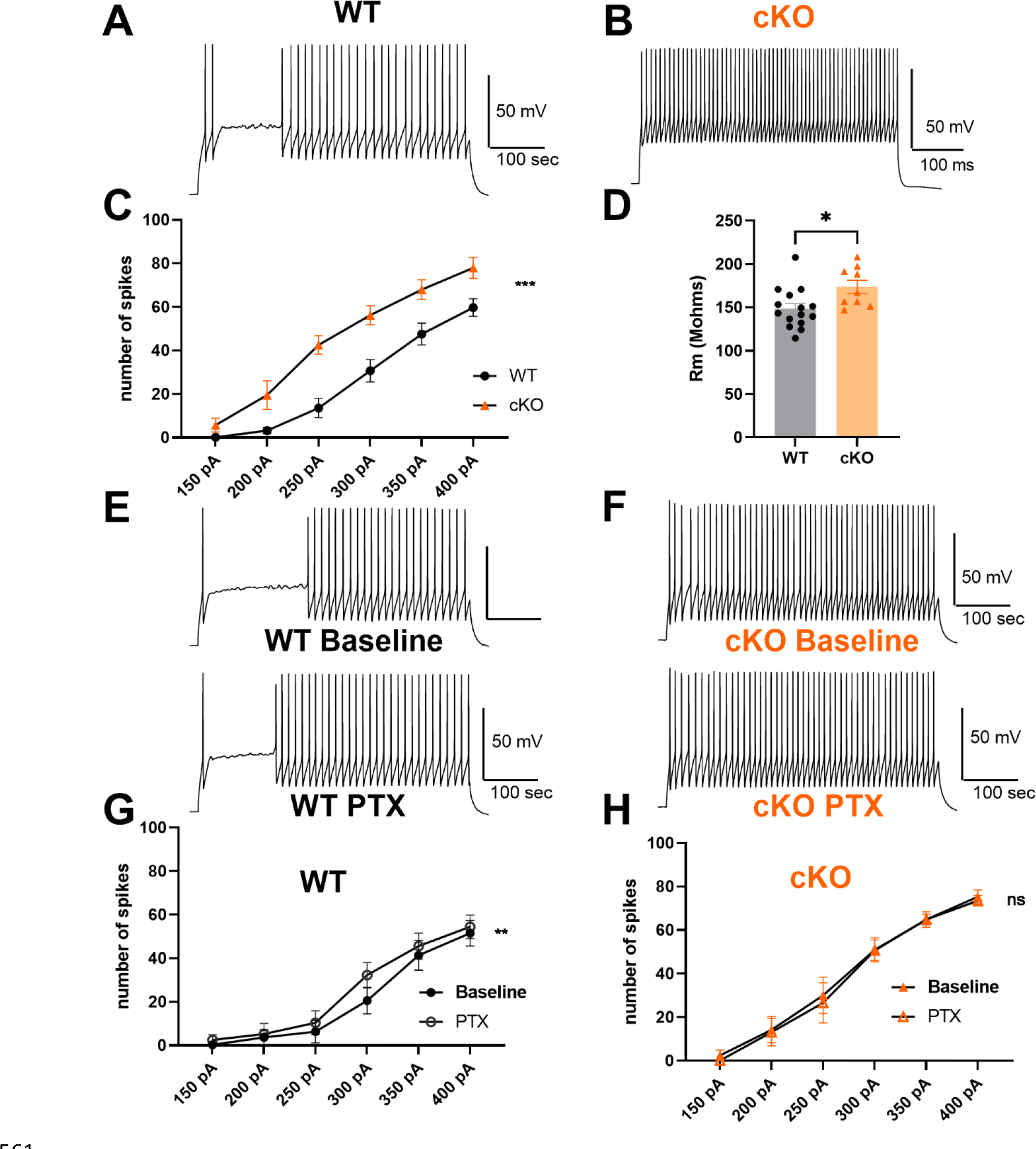
**cKO PV interneurons showed altered neuronal excitability from WT in mPFC. *A, B,*** Representative action potential patterns of WT (A) and cKO (B) PV interneurons with a 500 ms, 350 pA depolarizing current injection. ***C,*** Average number of action potentials in WT (N = 15) and cKO (N = 9) PV interneurons elicited over 500 ms at the indicated current amplitudes. Two-way ANOVA (F(1,22) = 14.43, P = 0.001) indicated a difference between WT and cKO PV interneurons. ***D,*** Summary of membrane input resistance for WT and cKO PV interneurons. Unpaired *t* test revealed differences between groups (p = 0.016). ***E, F,*** Representative action potential patterns of WT baseline, WT PTX, cKO baseline, and cKO PTX. 350 pA current was injected to elicit action potentials. ***G,*** Average number of action potentials in WT PV interneurons (baseline vs. 100 µM PTX, N = 11) elicited at the indicated current amplitudes. Two-way ANOVA showed a drug effect on number of action potentials (F(1, 10) = 15.44, P = 0.003). ***H,*** Average number of action potentials in cKO PV interneurons (baseline vs.

**Figure 2.**
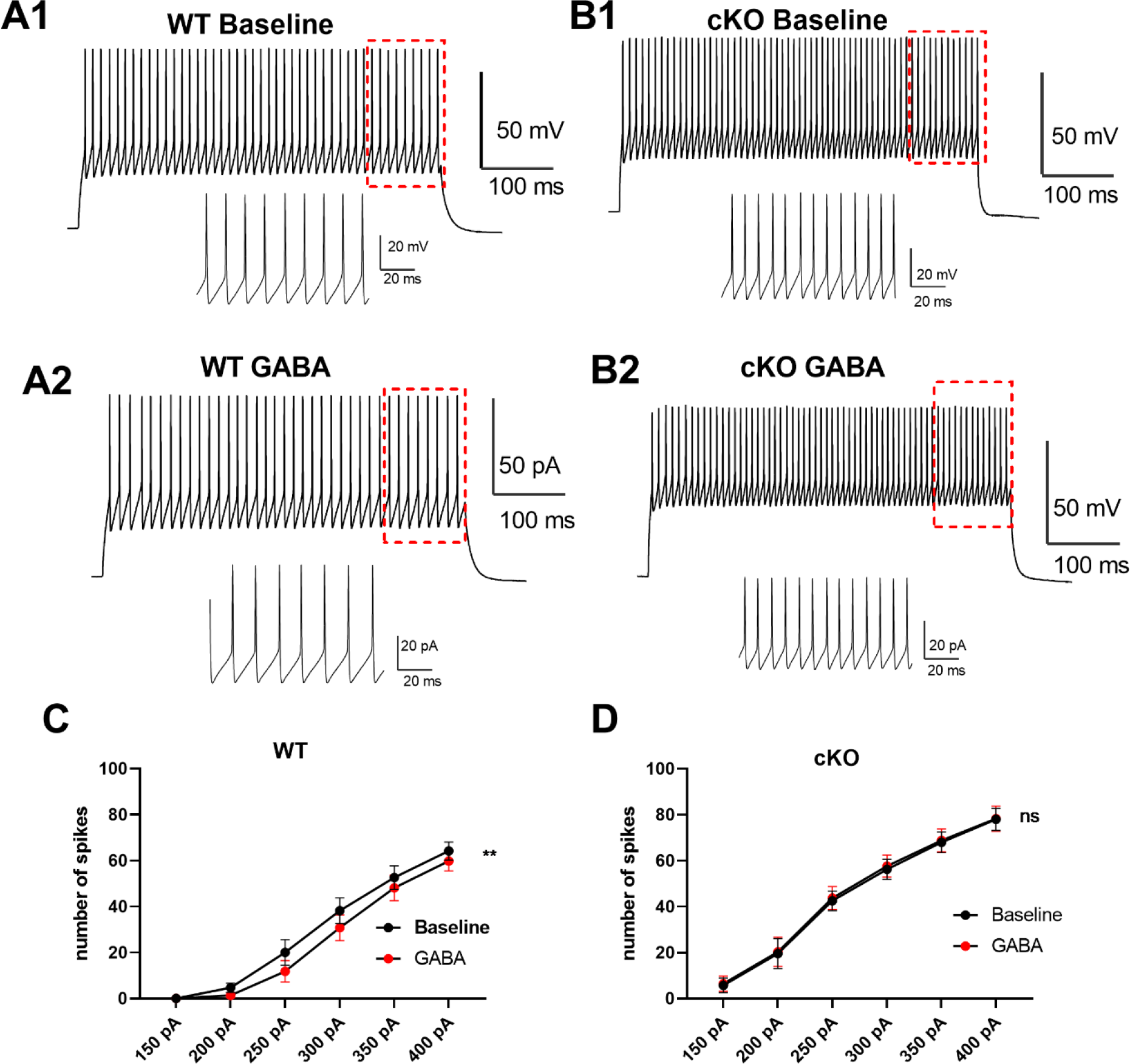
**GABA reduced action potentials in WT but not cKO PV interneurons. *A, B,*** Representative action potential patterns of WT baseline (A1), WT GABA (A2), cKO baseline (B1), and cKO GABA (B2). The red dashed box indicates the region highlighted in the inset, which shows individual action potentials at higher temporal resolution. ***C,*** Average number of action potentials in WT PV interneurons (baseline vs. 5 µM GABA, N = 10) elicited at the indicated current amplitudes. Two-way ANOVA showed a drug effect on number of action potentials (F(1, 9) = 13.26, P = 0.005). ***D,*** Average number of action potentials in cKO PV interneurons (baseline vs. 5 µM GABA, N = 9) elicited at the indicated current amplitudes. Two-way ANOVA showed no drug effect on number of action potentials (F(1, 8) = 0.494, P = 0.502).

**Figure 3.**
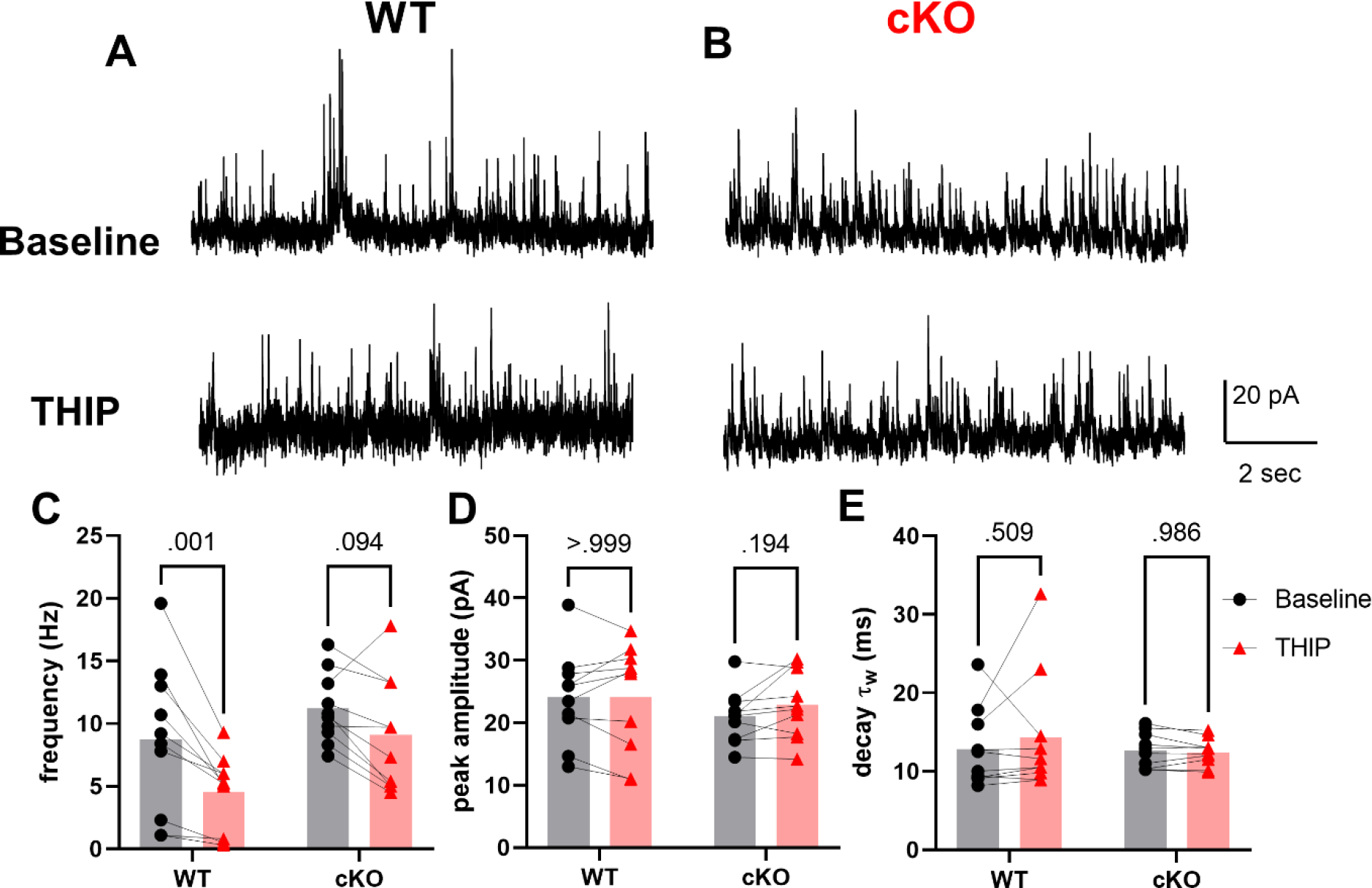
**THIP decreased the frequency of sIPSC in WT pyramidal neuron but not in cKO pyramidal neurons. *A, B,*** Representative WT pyramidal neuron (A) and cKO pyramidal neuron (B) held at 0 mV at baseline (top) or 1 µM THIP (bottom). ***C-E,*** sIPSC characteristics of WT pyramidal neurons (N = 10) and cKO pyramidal neurons (N = 10). ***C,*** Frequency of sIPSCs. Two-way ANOVA indicated a THIP effect (F(1,18) = 20.19, P < 0.001) but not genotype difference (F(1,18) = 3.872, P = 0.065), or an interaction between genotype and THIP (F(1,18) = 2.229, P = 0.153). Within each genotype, Sidak’s post-hoc test showed that THIP reduced the frequency of sIPSC in WT pyramidal neuron (P = 0.001) but not in cKO pyramidal neurons (P = 0.094). ***D,*** Peak amplitude of sIPSC. Two- way ANOVA showed no THIP effect (F(1,18) = 1.448, P = 0.244), genotype difference (F(1,18) = 0.529, P = 0.476), or interaction between genotype and THIP (F(1,18) = 1.519, P = 0.234). Within each genotype, THIP had no effect on the peak amplitude of sIPSC in WT and cKO (Sidak’s test, P > 0.999 and P = 0.194, respectively). ***E,*** weighted decay τ (τw) of sIPSC. Two-way ANOVA showed no THIP effect (F(1,18) = 0.419, P = 0.526), genotype difference (F(1,18) = 0.295, P = 0.594), or interaction between genotype and THIP (F(1,18) = 0.745, P = 0.399). Within each genotype, THIP had no effect on the τw of sIPSC in WT and cKO (Sidak’s test, P = 0.509 and 0.986, respectively).

In some experiments, we pharmacologically isolated sIPSCs of PV interneurons by maintaining a voltage of -70 mV with a whole-cell pipette solution of cesium chloride (in mM: 130 CsCl, 10 HEPES, 5 EGTA, 2 MgATP, 0.5 NaGTP, and 4 QX-314; pH adjusted to 7.3 with CsOH; 290 mOsm). The selective NMDA receptor antagonists NBQX (10 μM, Tocris Bioscience) and D-APV (50 μM, Tocris Bioscience) were added in the aCSF to inhibit ionotropic glutamate receptors. All phasic current recordings were performed in gap-free mode at 5 kHz sampling rate and 2 kHz filter cutoff frequency using an eight-pole Bessel filter.

### Measurement of kainate-induced action potentials

To induce network activity, 400 nM kainate in aCSF was perfused to slices. Cell-attached recordings were conducted using a pipette filled with aCSF. The number of action potentials before application of kainate and 10 min after kainate was measured and evaluated. Cell-attached recordings were performed in gap-free mode. The Chi square test was performed to compare the fraction of neurons exhibiting action potentials between WT and cKO.

### Measurement of local field potentials

As a measure of local-circuit activity, local field potential (LFP) recordings were conducted on 400 µm brain slices. Pipettes were filled with aCSF and placed at 100 µm depth in mPFC layer 2/3. LFP recordings were conducted in gap-free mode with sampling rate at 10 kHz and filter cutoff frequency between 2 Hz to 1 kHz using a bandpass filter. After 5 min baseline recordings, 400 nM kainate in aCSF was perfused for 10 min. The LFP recordings during the one-minute period before application of kainate and 10^th^ minute after kainate were used for analysis. The power spectral density of the signal was estimated using the *pspectrum* function in MATLAB (version R2023a)

### Bulk RNA-seq and differential expression analysis

Mice (3 WT and 3 cKO in each of two cohorts, for a total of 12 mice) were deeply anesthetized with isoflurane until unresponsive to tail pinch. Brains were rapidly removed, and tissue samples of frontal cortex were obtained and frozen at −80 °C until use. Total RNA was isolated from cortex tissue using RNeasy Plus Mini Kit (QIAGEN GmbH) in accordance with the manufacturer’s protocols. The integrity of total RNA was validated by an Agilent bioanalyzer. RNA samples that passed quality control were analyzed for sequencing by the Genome Technology Access Center at the McDonnell Genome Institute, Washington University School of Medicine. RNA samples were prepared according to the library kit manufacturer’s protocol, indexed, pooled, and sequenced on an Illumina NovaSeq X Plus. Basecalls and demultiplexing were performed using Illumina’s DRAGEN and BCLconvert version 4.2.4 software. RNA-seq reads were aligned to the Ensembl release 101 primary assembly using STAR version 2.7.9a (Dobin et al., 2013). Gene counts were obtained using Subread:featureCount version 2.0.3 (Liao et al., 2014). The ribosomal fraction, known junction saturation, and read distribution over known gene models were quantified using RSeQC version 4.0 (Wang et al., 2012).

Gene counts were imported into R/Bioconductor package EdgeR (Robinson et al., 2010) and TMM normalization size factors were calculated to adjust for differences in library size. Ribosomal genes and genes not expressed in at least one sample were excluded from further analysis. The TMM size factors and count matrix were then imported into R/Bioconductor package Limma (Ritchie et al., 2015). Weighted likelihoods based on the observed mean-variance relationship of every gene and sample were calculated for all samples with Limma’s voomWithQualityWeights (Liu et al., 2015). Differential expression analysis was performed to analyze differences between conditions. Results were filtered for genes with false discovery rate ≤ 0.05.

### Experimental design and statistical analyses

The number of action potentials was measured in Clampfit using peak detection algorithms. The membrane resistance was determined by measuring the steady-state voltage change in response to a −50 pA current. sIPSCs and sEPSCs were detected and analyzed as previously described using Clampfit algorithms (Lu et al., 2023). Data were analyzed and graphed using GraphPad Prism (version 9.0.0 for Windows, GraphPad Software). Paired *t* test or ANOVA as appropriate was conducted with Graphpad Prism.

## Results

### Deleting δ-containing GABAA receptors augmented PV interneuron excitability

In the central nervous system, δ-containing GABAA receptors primarily mediate tonic inhibition (Stell et al., 2003; Glykys et al., 2008; Lu et al., 2020). THIP, a δ-receptor preferring agonist, induces tonic currents in principal neurons of mouse hippocampus and neocortex (Drasbek and Jensen, 2006). Previously, we demonstrated that THIP elicited tonic currents in WT mPFC PV interneurons, predominantly mediated by δ- containing GABAA receptors (Lambert et al., 2024). Considering that genetic removal of δ receptors diminished tonic inhibition in response to the δ-preferring agonist, we hypothesized that the absence of δ-containing receptors might elevate the excitability of PV interneurons. We conducted current-clamp recordings and quantified the number of action potentials in response to a series of step-current injections in both WT and cKO PV interneurons (Fig. 1A, B). Notably, cKO PV interneurons exhibited significantly more action potentials compared to WT PV interneurons (Fig. 1C). Additionally, the input resistance of cKO PV interneurons was significantly elevated compared to that of WT PV interneurons (Fig. 1D). Consistent with a role for GABAA receptors in this altered excitability, 100 μM PTX increased firing in response to current injection in WT PV interneurons (Fig. 1E, G) but not in cKO PV interneurons (Fig. 1F, H). These results suggest a role for GABAA conductance in the excitability difference between WT and cKO PV interneurons.

If δ-containing receptors participate in PV interneuron excitability, we expect a difference in responsiveness of WT and cKO PV interneurons to the natural transmitter GABA, in addition to the previously demonstrated differential response to PTX (Fig.1).

In WT PV interneurons, 5 µM GABA decreased number of spikes in response to depolarizing current injection (Fig. 2A, C). On the contrary, GABA did not alter the number of action potentials in cKO PV interneurons (Fig. 2B, D). Surprisingly, these changes occurred in the absence of a detectable decrease in the input resistance at -70 mV in either genotype, suggesting excitability measures are more sensitive (100 ± 8% change for WT, N = 10; 103 ± 6% change for cKO, N = 9).

GABAergic PV interneurons release the inhibitory neurotransmitter GABA onto pyramidal neurons (Hu et al., 2014). Therefore, altering the excitability of PV interneurons might be expected to alter transmitter release detected by layer 2/3 pyramidal neurons. To test this, we applied 1 µM THIP to slices while recording sIPSCs in WT and cKO pyramidal neurons (Fig. 3A, B). Although we did not detect a significant interaction between genotype and drug, post hoc Sidak tests showed that THIP decreased the frequency of sIPSCs in WT but not pyramidal neurons from PV-cKO slices (Fig. 3C). Neither the peak amplitude nor the kinetics of sIPSCs was altered by THIP in WT or cKO recordings (Fig. 3D, E). In line with previous findings (Drasbek et al., 2007), THIP also altered the excitatory/inhibitory (E/I) ratio of WT layer 2/3 pyramidal neurons by reducing the frequency of sIPSCs while increasing the frequency of sEPSCs (extended Fig. 3-1).

We reasoned that PV interneurons themselves receive input from non-PV interneurons, which are not δ bearing and thus should not be affected by THIP application. Indeed, THIP had no effect on the frequency or decay of sIPSCs in PV interneurons of either WT or cKO slices (Fig. 4). THIP did increase the amplitude of sIPSCs in WT PV interneurons (Fig. 4D), which could be caused by the increase of noise of baseline masking sIPSCs with small amplitude. The effect on amplitude was not observed in cKO cells.

**Figure 4.**
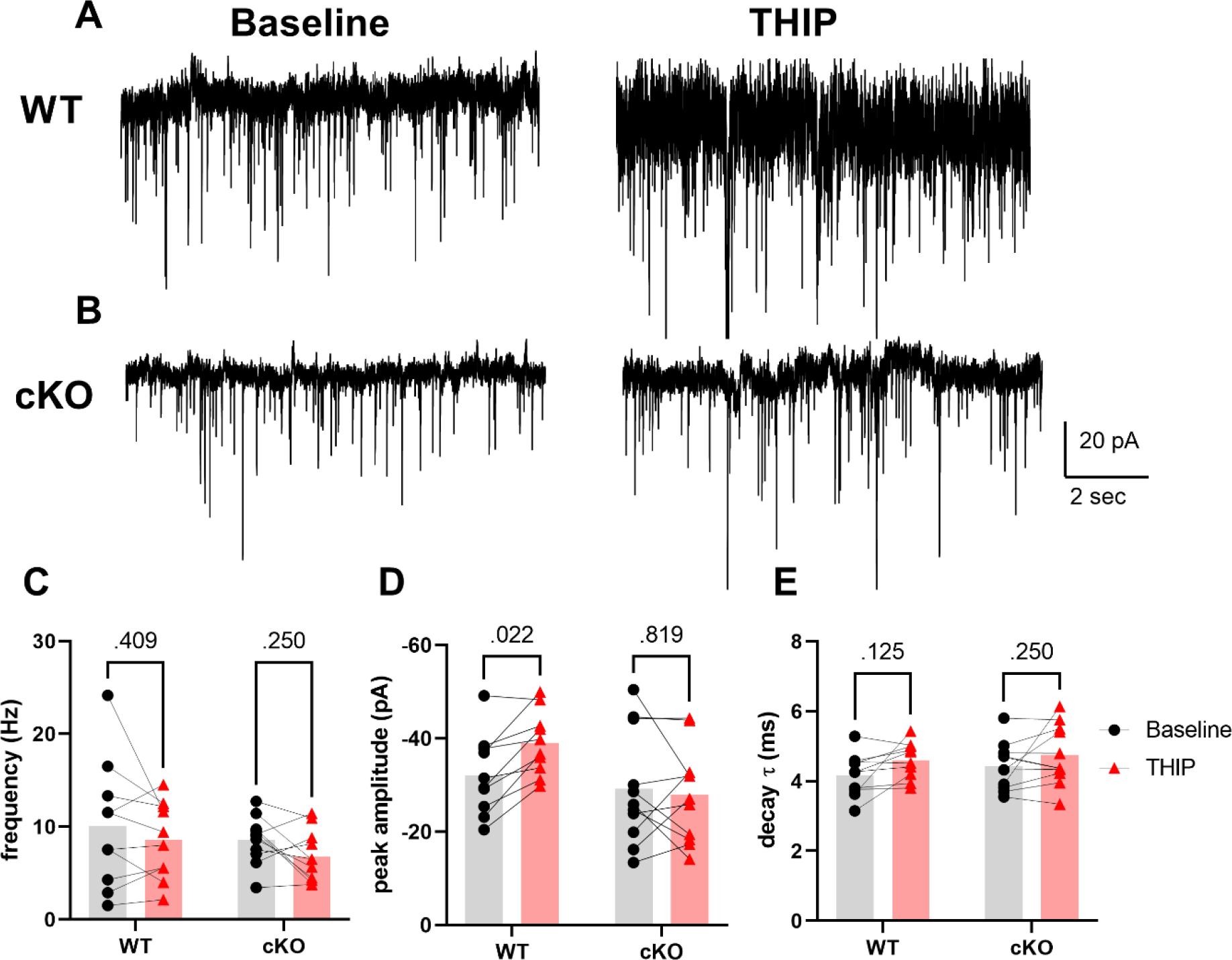
**THIP had no effect on the frequency of sIPSC in WT or cKO PV interneurons. *A,*** Representative trace of a WT PV interneuron held at -70 mV at baseline (left) or in the presence of 1 µM THIP (right). ***B,*** Representative traces of a cKO PV interneuron held at -70 mV at baseline (left) or in the presence of 1 µM THIP (right). ***C-E,*** sIPSC characteristics of WT PV interneurons (N = 10) and cKO PV interneurons (N = 11). ***C,*** frequency of sIPSC. Two-way ANOVA showed no THIP effect (F(1,19) = 3.9, P = 0.063), genotype difference (F(1,19) = 0.935, P = 0.346), or interaction between genotype and THIP (F(1,19) = 0.034, P = 0.855). Within each genotype, THIP did not alter the frequency of sIPSC in WT or cKO PV interneurons (Sidak’s test, P = 0.409 and 0.25, respectively). ***D,*** Peak amplitude of sIPSC. Two-way ANOVA showed no THIP effect (F(1,19) = 2.707, P = 0.116), or genotype difference (F(1,19) = 3.181, P = 0.090). There is an interaction between genotype and THIP (F(1,19) = 5.928, P = 0.025). THIP increased the peak amplitude of sIPSC in WT PV interneurons but not cKO PV interneurons (Sidak’s test, P = 0.022 and 0.819, respectively). ***E,*** weighted decay τ (τw) of sIPSC. Two-way ANOVA showed no genotype difference (F(1,19) = 0.563, P = 0.462), or interaction between genotype and THIP (F(1,19) = 0.115, P = 0.738). THIP increased the τw throughout genotypes (F(1,19) = 6.247, P = 0.022). Within each genotype, THIP had no effect on the τw of sIPSC in WT and cKO (Sidak’s test, P = 0.125 and 0.25, respectively).

### Deleting δ subunits in PV interneurons alters the excitability of pyramidal neurons and network activity

To investigate whether the altered phasic inhibition of pyramidal neurons would impact the excitability of pyramidal neurons, we applied a low concentration (400 nM) of kainate to excite the network. Kainate activates kainate receptors, a subset of ionotropic glutamate receptors (Hadzic et al., 2017) but also is a non-desensitizing AMPA receptor agonist (Paternain et al., 1995) and may activate a small percentage of these receptors (Cherubini et al., 1990; Jiang et al., 2001). Indeed, 400 nM kainate induced steady tonic current, sensitive to 10 µM NBQX, in layer 2/3 pyramidal neurons (0.436 ± 0.134 pA/pF, N = 6) and in PV interneurons (0.447 ± 0.106 pA/pF, N = 4). In cell-attached recordings of pyramidal neurons, kainate induced robust firing in WT but not in cKO pyramidal neurons (Fig. 5A-C).

**Figure 5.**
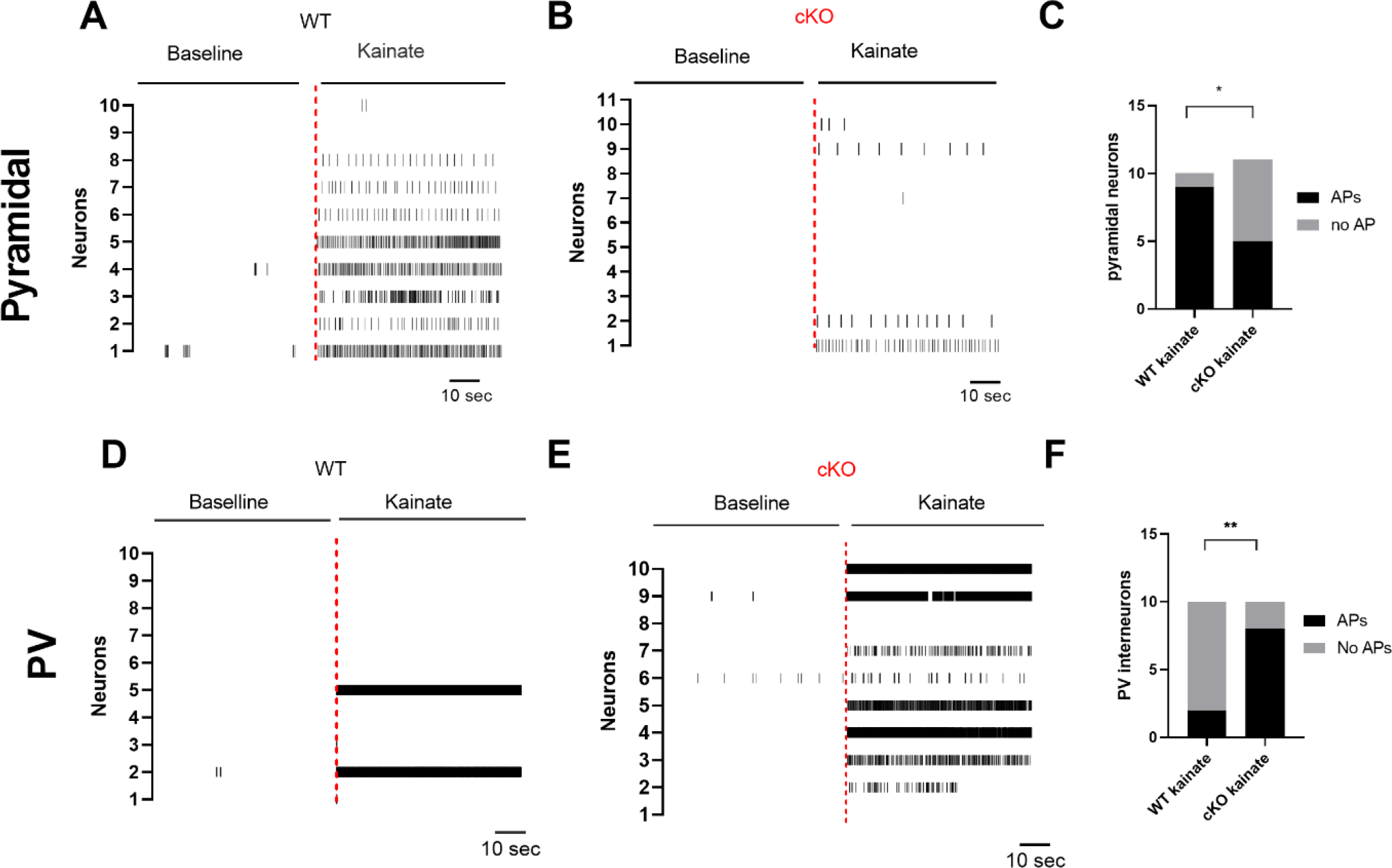
**Kainate-induced neuronal excitability was altered in cKO. *A, B,*** Spike raster plot of pyramidal neuron firing in response to kainate application, compared to baseline, in WT (A) and cKO (B) mice. ***C,*** Quantification of the fraction of pyramidal neurons exhibiting action potentials (APs) or no APs during kainate application in WT and cKO conditions. More pyramidal neurons exhibited APs in the WT condition upon kainate treatment (chi square test, P = 0.031). ***D, E,*** Spike raster plot of PV interneuron firing in response to kainate application, compared to baseline, in WT (D) and cKO (E) mice. ***F,*** Fraction of PV interneurons exhibiting APs or no APs during kainate application in WT and cKO conditions. More PV interneurons exhibited APs in the cKO condition upon kainate treatment (chi square test, P = 0.007).

We hypothesized that the reduction of kainate effects on cKO pyramidal neurons may result from the higher excitability of cKO PV interneurons and increased GABA output to these principal neurons (Figs. 1-3). To test this, we recorded kainate-induced action potentials from cell-attached recordings of PV interneurons (Fig. 5D, E) and observed reciprocal effects compared with pyramidal neurons; WT PV cells showed little responsiveness to kainate, but cKO PV neurons exhibited strong responsivity to kainate. These results demonstrate the strong influence of δ-containing GABAA on two major cell types of the cortex.

We explored the implications of these excitability differences on local network function. In the mPFC, PV interneurons play a crucial role in generating brain oscillations. To investigate how deletion of δ-containing receptors would affect oscillations, we conducted local field recordings in mPFC layer 2/3 during application of kainate.

Although 400 nM kainate has previously been shown to selectively increase gamma oscillations (Gloveli et al., 2005), our results showed that kainate induced broad oscillations in WT slices (Fig. 6A,B, E). In contrast, kainate failed to induce broadband power increases in cKO cortex (Fig. 6C, D, E). These results show that that a single receptor class in one group of interneurons has a strong impact on overall network function.

**Figure 6.**
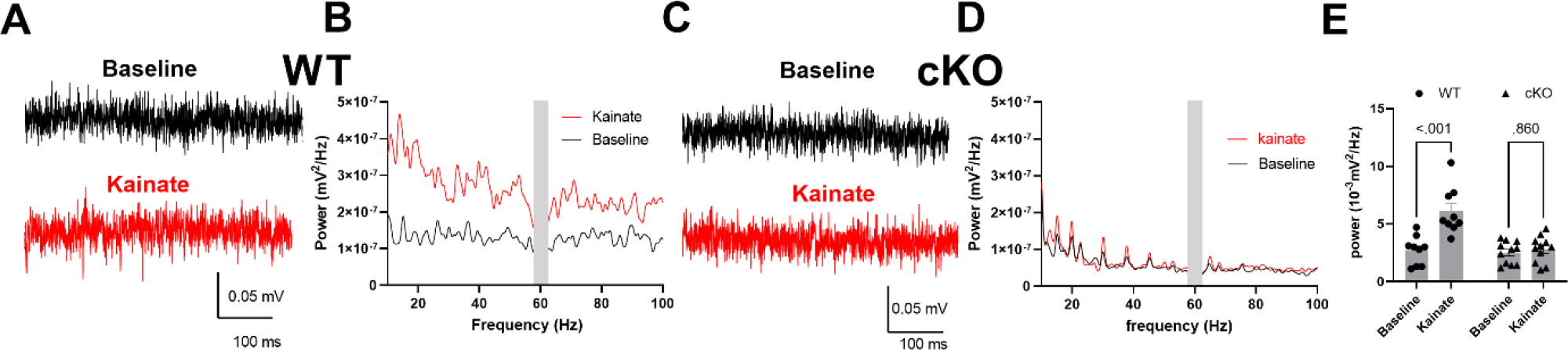
**Kainate induced broadband oscillations in mPFC layer 2/3 of WT but not cKO mice. *A,*** Representative LFP traces recording from mPFC layer 2/3 in a WT mouse slice at baseline (top) and 10 min after application of kainate (bottom). ***B,*** Power spectral density plot showing the frequency composition of the LFP signal at baseline (black) and after kainate application (red) in the WT slice. Kainate induced a broadband increase in oscillatory power across multiple frequencies. ***C,*** Representative LFP traces from a cKO mouse slice at baseline (top) and 10 min after kainate application (bottom). ***D,*** Power spectral density plot for the cKO slice, showing little change in oscillatory power with kainate application compared to baseline across most frequencies. ***E***, Quantification of oscillatory power from 15 – 80 Hz during baseline and kainate application in WT and cKO slices. Two-way ANOVA showed an interaction between genotype and kainate (F(1,18) = 12.56, P = 0.002), as well as differences on genotypes (F(1,18) = 15.76, P < 0.001) and kainate effect (F(1,18) = 17.72, P <0.001). Post- hoc tests showed that kainate increased the broadband power in WT (Sidak’s test, P < 0.001) but not cKO slices (Sidak’s test, P = 0.86). The vertical gray bars in panel B and D indicate the frequency range (59-61 Hz) that was notch-filtered to remove 60 Hz noise.

Our findings demonstrate that deletion of δ-containing GABAA receptors in PV interneurons alters excitability of both PV interneurons and pyramidal neurons through acute effects on GABA inhibition, revealed by antagonist and agonist application.

Beyond direct impact on GABAergic inhibition in the cKO model, we sought to preliminarily investigate whether deletion of δ subunits in PV interneurons influences cortical gene expression, potentially contributing to observed excitability differences. To address this, we performed bulk RNA-seq analysis on six mice in each genotype. Based on a false discovery rate criterion of 0.05 and expression difference cutoff of 2-fold, no genes were differentially expressed between the WT and cKO groups. Extended Table 6-1 shows the detailed results for top transcripts. Full results are available at Figshare (https://doi.org/10.6084/m9.figshare.26026741.v1). These hypothesis-generating results do not allow us to exclude conclusively a contribution of secondary transcriptional changes to excitability differences. Nevertheless, no immediate gene candidates emerged to explain cortical excitability differences in response to the cKO manipulation.

## Discussion

In this study, we investigated the role of δ-containing GABAA receptors in PV interneurons of the mPFC and their impact on neuronal excitability, network oscillations, and overall circuit function. Using a *Gabrd* cKO mouse line, we uncovered a crucial role for these receptors in regulating the excitability of PV interneurons and, consequently, the inhibition exerted by these interneurons on pyramidal neurons. The impact of these specialized receptors has been unclear, especially given the prominence of more conventional GABAA receptors (e.g., those containing a γ2 subunit) at synapses (Sieghart and Sperk, 2002), demonstrated herein by the lack of impact of δ loss on phasic inhibition in PV interneurons (Figure 4).

Our results demonstrate that the absence of δ-containing GABAA receptors leads to increased excitability of PV interneurons, as evidenced by their heightened firing rates and elevated input resistance compared to WT PV interneurons. This increased excitability can be attributed to the loss of tonic inhibition mediated by δ-containing GABAA receptors, which normally dampens the excitability of PV interneurons. The observed effects of the GABAA receptor antagonist PTX and the δ-preferring agonist THIP further support the involvement of these receptors in regulating PV interneuron excitability. The strong impact of the loss of δ is perhaps surprising, as it is evident from analysis of sIPSCs that strong phasic inhibition persists in PV interneurons when δ is missing in these cells (e.g., Figure 4, (Lambert et al., 2024)).

Notably, the increased excitability of PV interneurons in the cKO mice had profound consequences for the inhibitory control exerted by these interneurons on pyramidal neurons. The frequency of sIPSCs in WT pyramidal neurons was reduced upon THIP application, indicating decreased release of GABA from PV interneurons. Conversely, THIP failed to alter sIPSC frequency in cKO pyramidal neurons, likely due to the heightened excitability and aberrant inhibitory output of PV interneurons lacking δ- containing GABAA receptors.

The altered excitability of PV interneurons and the consequent dysregulation of inhibitory control over pyramidal neurons had significant implications for overall network dynamics in the mPFC. We found that kainate-induced oscillations, which are known to depend on the activity of PV interneurons (Fuchs et al., 2007; Ferando and Mody, 2013), were reduced in the cKO cortex compared to WT slices. This observation highlights the critical role of δ-containing GABAA receptors in shaping the oscillatory activity of the mPFC, which is crucial for coordinating neuronal activity and facilitating communication between brain regions during cognitive processes. For these observations, we acknowledge the limitations of the preparation chosen. Although the *ex vivo* slice preparation allowed us to draw conclusions about intrinsic cortical network behavior free of subcortical influences and long-range cortical interactions, it is possible that the results would differ in the intact animal because of these extrinsic influences.

We reasoned that altering GABAergic inhibition in PV interneurons over the course of development could result in transcriptional differences that could participate in excitability changes. As a preliminary approach, we performed an unbiased bulk transcriptional analysis of cortex. Our analysis, meant to detect broad transcriptional changes that might affect overall excitability evident in Figures 5 and 6, did not reveal any genes differentially expressed in cKO cortex. It is possible, as hypothesized, that the deletion of δ-containing GABAA receptors in PV interneurons primarily affects their functional properties and connectivity without inducing substantial transcriptional changes. Alternatively, the bulk approach used in this study may have missed cell type-specific transcriptional changes. PV interneurons constitute a small proportion of the total cell population in the cortex, so cell-autonomous transcriptional changes may be missed in favor of transcriptional profiles of other cell types in the bulk tissue sample. To address these limitations, future studies could employ single-cell RNA-seq techniques (Hwang et al., 2018), which enable the profiling of gene expression at the individual cell level and allow for the identification of cell-type specific transcriptional changes.

The findings of this study have important implications for understanding neural mechanisms underlying cognitive functions mediated by the mPFC, such as working memory, attention, and decision-making. Dysregulation of PV interneuron function and disruption of mPFC network dynamics have been implicated in various neuropsychiatric disorders, including schizophrenia and autism spectrum disorders (Xu et al., 2019). Our results provide insights into the specific contribution of δ-containing GABAA receptors in shaping the excitability and inhibitory output of PV interneurons in mPFC, offering potential avenues for therapeutic interventions targeting these receptors or the associated signaling pathways (Thompson, 2024).

Although our study utilized acute slice preparations, future research should explore the impact of chronic alterations in δ-containing GABAA receptor expression on PV interneuron function and mPFC network dynamics in vivo. Additionally, behavioral studies beyond those performed (Lambert et al., 2024) should investigate the cognitive and behavioral consequences of selectively manipulating these receptors in PV interneurons to further elucidate their functional relevance. These extensions of the present work will be important to rational development of drugs that target δ-containing GABAA receptors, of current interest neurospsychiatry (Whissell et al., 2015; Maguire and Mennerick, 2023; Thompson, 2024).

In conclusion, our findings highlight the pivotal role of δ-containing GABAA receptors in regulating the excitability and inhibitory output of PV interneurons in the mPFC, thereby shaping the balance between excitation and inhibition necessary for proper network function and cognitive processing. These insights contribute to our understanding of the neural circuits and mechanisms underlying mPFC-dependent cognitive functions and may guide future research into the pathophysiology and potential therapeutic targets for neuropsychiatric disorders associated with mPFC dysfunction.

**Conflict of Interest**. CFZ is a member of the Scientific Advisory Board for Sage Therapeutics and holds equity in Sage Therapeutics. Sage Therapeutics had no role in the design or interpretation of the experiments herein. The remaining authors declare no competing financial interests.

## Acknowledgements

The work was funded by NIMH grants R01 MH123748 and P50 MH122379 (SM and CFZ) and F30 MH126548 (PML). The authors thank members of the Taylor Family Institute for Innovative Psychiatric Research for discussion and input. We thank Dr. Jamie Maguire for a gift of the floxed δ mice and Dr. Yanbo Yu for preliminary help on differential gene expression.

## Author contributions

Conceptual study design: XL & SM. Data collection and analysis: XL, HS, & PL. Original draft: XL. Critical revisions: XL, CFZ, & SM.

---

**Extended Figure 3-1.**
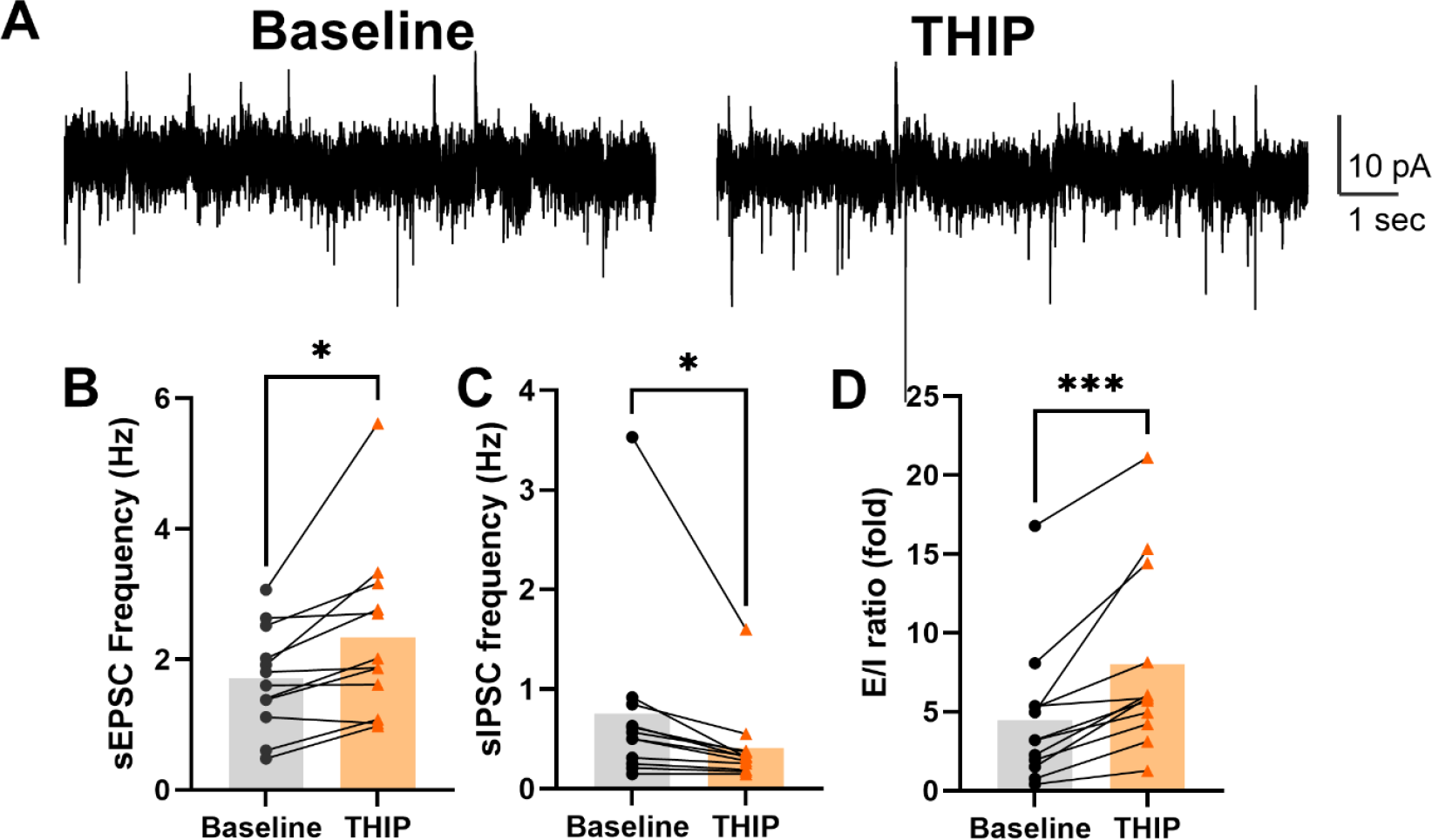
**THIP altered the E/I ratio in mPFC layer 2/3 pyramidal neurons. *A,*** Representative trace of a WT pyramidal neuron held at -40 mV at baseline (left) or in the presence of 1 µM THIP (right). Both sIPSCs (upward) and sEPSCs (downward) were recorded simultaneously. ***B,*** THIP increased the frequency of sEPSCs (paired t test, P = 0.013). ***C,*** THIP decreased the frequency of sIPSCs (paired t test, P = 0.045). D, THIP increases the E/I ratio (paired t test, P = 0.001).

**Extended Table 6-1.**
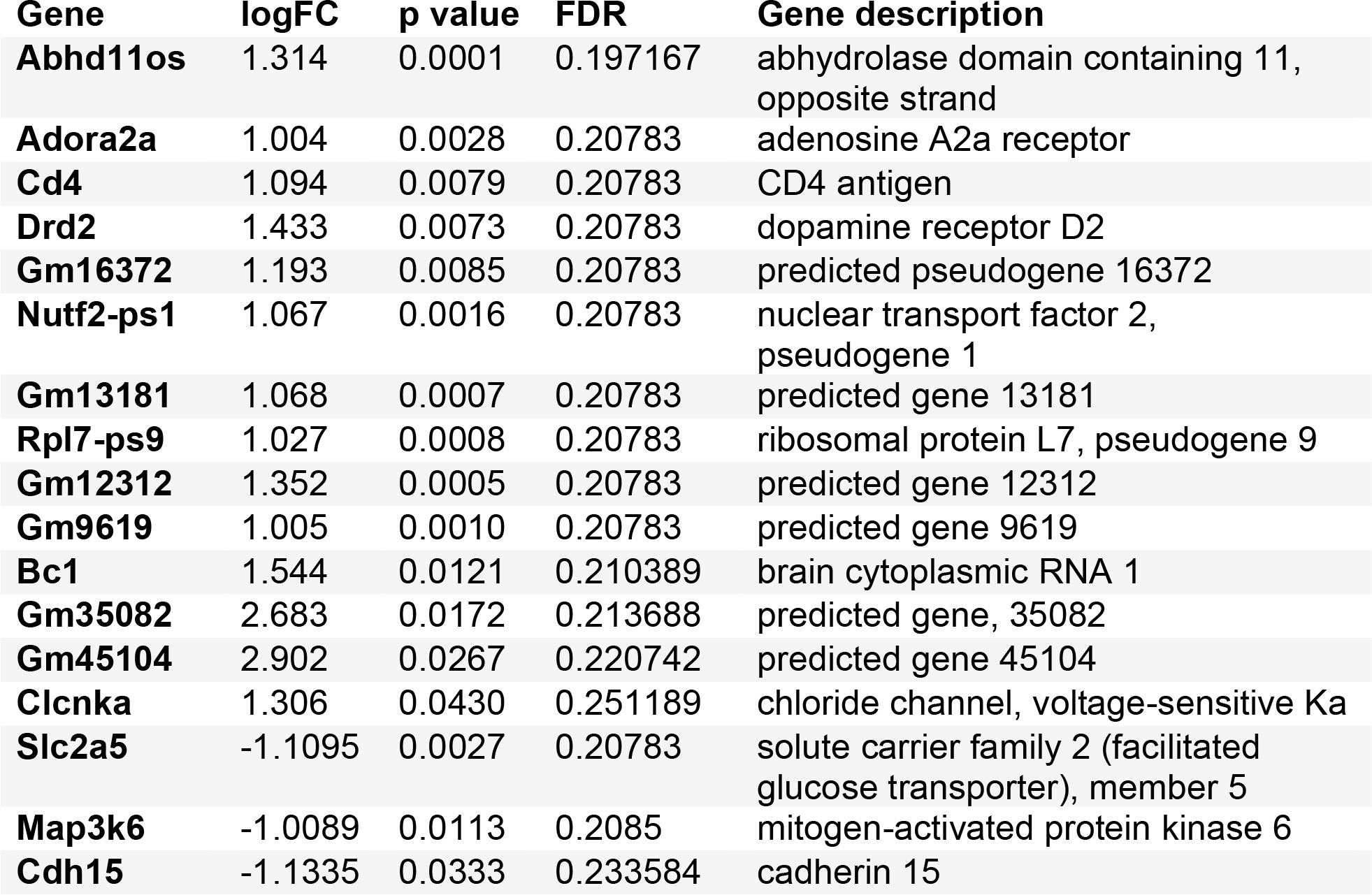
**List of candidates for differentially expressed genes in the cortex.** The log2 fold change (logFC), p values, and false discovery rate (FDR) are listed for transcripts that met criteria of logFC >1 (either direction) and an uncorrected p value of 0.05. A full list of transcripts is available at Figshare (https://doi.org/10.6084/m9.figshare.26026741.v1).

